# Defining the Neural Kinome: Strategies and Opportunities for Small Molecule Drug Discovery to Target Neurodegenerative Diseases

**DOI:** 10.1101/2020.04.01.020206

**Authors:** Andrea I. Krahn, Carrow Wells, David H. Drewry, Lenore K. Beitel, Thomas M. Durcan, Alison D. Axtman

**Author notes:** Corresponding Author, Alison D. Axtman – Structural Genomics Consortium, UNC Eshelman School of Pharmacy, University of North Carolina at Chapel Hill, Chapel Hill, NC, 27599, USA; Division of Chemical Biology and Medicinal Chemistry, UNC Eshelman School of Pharmacy, University of North Carolina at Chapel Hill, Chapel Hill, NC, 27599, USA.

## Abstract

Kinases are highly tractable drug targets that have reached unparalleled success in fields such as cancer but whose potential has not yet been realized in neuroscience. There are currently 55 approved small molecule kinase-targeting drugs, 48 of which have an anti-cancer indication. The intrinsic complexity linked to central nervous system (CNS) drug development and a lack of validated targets has hindered progress in developing kinase inhibitors for CNS disorders when compared to other therapeutic areas such as oncology. Identification and/or characterization of new kinases as potential drug targets for neurodegenerative diseases will create opportunities for development of CNS drugs in the future. The track record of kinase inhibitors in other disease indications supports the idea that with the best targets identified small molecule kinase modulators will become impactful therapeutics for neurodegenerative diseases.

**KEY CONCEPTS:** **Chemical probe**: a high-quality small molecule that is potent, selective, and cell-active that meets the following criteria: (1) *in vitro* biochemical IC_50_ < 50 nM, (2) ≥ 30-fold selectivity relative to other kinases in a large assay panel such as DiscoverX *scanMAX*, and (3) cellular activity or target engagement with an IC_50_ < 1 μM

**Narrow spectrum**: a selectivity threshold that can be defined as potently inhibiting ∼10% or less of all kinases screened

**Kinome**: all human kinases

**Kinase chemogenomic set (KCGS)**: publicly-available curated physical library of narrow spectrum and potent kinase inhibitors for which the SGC-UNC has received permission to share the compounds; subsequent releases will increase kinome-wide coverage

**Illuminating the Druggable Genome (IDG) program**: several interconnected projects currently funded by the National Institutes of Health to provide information on historically understudied members within protein families that have provided drug targets; the three main focus areas are kinases, G-protein coupled receptors, and ion channels

**IDG kinase**: a kinase that was nominated as dark (understudied) by the National Institutes of Health IDG program (curated list found here: https://druggablegenome.net/IDGProteinList); IDG consortium members generate data and resources to aid in the illumination of the function of these kinases

**DK tool**: a narrow spectrum inhibitor that exhibits a defined selectivity score (S_10_(1 μM) < 0.05) and cellular target engagement with an IC_50_ < 1 μM; S_10_(1 μM) is a measure of selectivity equal to the percentage of screened kinases biochemically inhibited by >90% at 1 μM

## 1. INTRODUCTION

There is an urgent and growing need for new drugs to treat chronic neurogenerative diseases, as life expectancies continue to increase and, correspondingly, the number of patients rises on an annual basis. Unfortunately, there is still much to be learned about the pathogenesis of these complex brain disorders and the ability to treat them is often hampered by the difficulties associated with diagnosis, leading to intervention too late. The heterogeneity of these diseases contributes to their complexity. Neurodegenerative disorders are often diverse syndromes with multiple subtypes and the one-size-fits-all treatment model, therefore, results in failure. Following the approach that has yielded clinical success in oncology, personalized medicine could similarly be applied to neurodegeneration. This situation is further complicated by the fact that the likelihood of U.S. Food and Drug Administration (FDA) approval for CNS drugs is approximately half that for drugs with a peripheral target and the approval process takes nearly twice as long as for an oncology drug.^*1*, *2*^ Finally, the failure of many advanced clinical trials in the area of Alzheimer’s disease (AD) and other diseases of the brain has caused many pharmaceutical companies to exit the neuroscience research and development space, reducing the resources being invested to address the problem.

As an impetus to drive CNS-based kinase drug discovery, we provide insight into the tractability of a subset of human kinases with links to neurodegeneration in this review. Rather than focusing on the few widely studied kinases around which most of the data, reagents, and small molecules have been generated, we instead concentrate on detailing the chemical tools and biology that will encourage and enable the study of lesser studied (also known as understudied or dark) kinases. This focus on the understudied kinome is aligned with that of the Structural Genomics Consortium (SGC), which aims to promote the exploration of currently understudied, but potentially disease-relevant proteins across the human proteome.^*3*^ In addition to characterizing which kinases fall into this understudied subcategory, we offer high-quality chemical starting points that can be used to modulate their activity. Furthermore, through data mining, we have tabulated genetic links between understudied kinases and the most common adult-onset neurogenerative diseases: AD, Parkinson’s disease (PD), and amyotrophic lateral sclerosis (ALS).

Since not all protein kinase targets can be covered in a single review, we provide an overview of promising understudied human protein kinase targets for AD, PD, and ALS. Furthermore, we focus our attention on tractable kinases for which there exists a small molecule with narrow selectivity profile against the screenable human kinome (i.e. a non-promiscuous kinase inhibitor). The Illuminating the Druggable Genome (IDG) program has identified many of these understudied kinases on their understudied protein list (https://druggablegenome.net/IDGProteinList). We conclude with a discussion of the challenges that remain with respect to small molecule development for validated biological targets in neurodegenerative disorders.

## 2. COMMON CHRONIC NEURODEGENERATIVE DISEASES

AD is the most common cause of dementia, accounting for between 60-80% of cases. An estimated 5.8 million Americans of all ages are living with AD in 2019.^*4*^ It is a disease that develops over many years, occurring in slowly progressive stages before symptoms manifest. The two primary hallmark pathologies of AD are amyloid-β (Aβ) plaques, which are the extracellular accumulation of the Aβ peptide, and neurofibrillary tangles (NFTs) composed of abnormally phosphorylated tau.^*5*^ These NFTs have been observed in the entorhinal cortex, the hippocampus, the amygdala, and the neocortex, with the degree of dementia correlating with the distribution of NFTs.^*6*^ Additionally, brain tissue from AD patients demonstrates the presence of activated microglia and astrocytes surrounding Aβ plaques, as well as elevated cytokine levels, indicative of inflammation as a contributing factor in the development of the disease.^*7*, *8*^

PD is the most common movement disorder and the second most common age-related neurodegenerative disorder after AD. An estimated one million Americans are thought to have PD, with nearly 60,000 newly diagnosed cases per year and thousands living undiagnosed.^*9*^ PD is characterized by the selective and progressive degeneration of dopamine (DA) producing neurons in an area deep in the brain known as the substantia nigra (SN).^*5*^ Loss of the neurotransmitter DA results in the classical clinical motor symptoms that include bradykinesia, rigidity, postural instability, and resting tremor. The remaining neurons in the SN typically contain proteinaceous inclusions, termed Lewy bodies, which have historically been thought to be mainly composed of alpha-synuclein and ubiquitin.^*10*^ A recent study, however, demonstrated that Lewy bodies are composed of a membranous medley rather than protein fibrils, which includes membrane fragments, lipids, and other cellular material.^*11*^ More recently, studying immune dysregulation in PD has attracted increased interest as activated microglia and T cells have been found in the SN of PD patients.^*12*^ Aside from the motor symptoms of PD, patients exhibit non-motor features years before clinical diagnosis, including a loss of smell, depression, anxiety, constipation, and the biggest predictor of PD in prodromal patients, rapid eye movement (REM) sleep behavior disorder (RBD).^*10*^ RBD is characterized by loss of REM sleep paralysis, allowing patients to “act out” dreams and represents the highest predictive value for PD, giving clinicians the ability to intervene early in the course of the disease before neuronal loss progresses past rescue and before symptomatic therapies confound assessments.^*13*^

ALS is the most common motor neuron (MN) disorder and the third most common adult-onset neurodegenerative disease after AD and PD. The ALS Association estimates the current prevalence of ALS is between 12,800 and 19,800 people in the United States.^*14*^ ALS is a progressive neurological disorder that affects both upper MNs (projecting from the cerebral cortex to the spinal cord) and lower MNs (projecting from the spinal cord to muscles). It is pathologically characterized by cytoplasmic inclusions of MNs primarily composed of ubiquitinated TDP-43 protein, except for patients with SOD1 or FUS mutations, in which case the main component of the inclusion bodies are SOD1 protein and FUS protein, respectively.^*15*^ Clinical features of the disease are mainly associated with motor dysfunction (muscle weakness, spasticity, and atrophy), but almost half of patients also develop cognitive and/or behavioral symptoms.^*15*, *16*^ Neuroinflammation has also been implicated in ALS as activated microglia and astrocytes are present in post-mortem tissue, and altered cytokine profiles have been reported.^*17*^

## 3. CURRENT DRUG LANDSCAPE AND KINASES AS THE FUTURE

Currently available drugs to treat AD, PD, and ALS target only symptoms and do not offer substantial benefit to patients, having modest effects at best. Only two types of medications have been approved by the FDA for treating cognitive symptoms of AD: cholinesterase inhibitors and the N-methyl-D-aspartate receptor agonist memantine. To date, five drugs have been approved for the treatment of AD, and since 2003 no new drugs have been approved, with most failed drugs focused on targeting Aβ accumulation.^*18*^ Late in 2019 Biogen announced that they were applying for FDA marketing approval of their anti-amyloid human monoclonal antibody known as aducanumab. Aducanumab, which selectively binds to Aβ fibrils and soluble oligomers, demonstrated substantial reduction of Aβ plaques in a dose- and time-dependent manner, such that after 12 months nearly half the patients who received the 10 mg/kg dose no longer had positive amyloid PET scans.^*19*^

Levodopa (L-DOPA), a precursor of DA, has been the most effective drug for treating the motor symptoms of PD but has increasingly negative side effects as the disease progresses. Other FDA approved treatments for PD are predominantly focused on increasing or substituting for low levels of DA to manage tremors and irregular movements: DA agonists directly stimulate the DA receptors, monoamine oxidase B inhibitors prolong DA stimulation by decreasing DA breakdown, and catechol-O-methyl transferase inhibitors delay the breakdown of L-DOPA.^*10*^ Non-dopaminergic treatments include anticholinergics, adenosine 2A antagonists, and a drug that is reported to act via multiple mechanisms to exert anti-Parkinsonian effects (amantadine).^*20*^ These FDA-approved drugs can be used as alternatives to L-DOPA, in addition to L-DOPA, or in combination with one another. Patients with advanced PD, who are unresponsive to drug treatment can alternatively be treated with deep brain stimulation to control motor symptoms.^*21*^

Treatments for ALS focus on managing symptoms through physiotherapy, nutrition, and respiratory support. The three FDA approved medications, two of which are different formulations of the same active agent, are only mildly effective at prolonging survival and are often taken in combination with symptomatic treatments. Since the first approved drug was hypothesized to modulate glutamate transmission, a vast majority of compounds tested in the past couple of decades have been anti-glutamatergic but have not been successful. Other attempts have been made to reduce inflammation and oxidation, but most compounds have failed to demonstrate efficacy in human trials.^*22*^

In spite of the ability of the biomedical research community to turn kinase inhibitors into approved medicines, we believe this target tractability has not been exploited to the degree possible in neuroscience. By January 2020, the FDA had approved 55 small molecule protein kinase inhibitors and 53 of them are orally effective. The data collected around those 55 supports that 48 of them are used in the treatment of neoplastic diseases, while 9 are employed in the treatment of non-malignancies such as glaucoma and rheumatoid arthritis, for example. Remarkably, 17 have found utility in the treatment of more than one disease, such as imatinib, which is approved for the treatment of eight disparate disorders.^*23*^ Kinases have been largely ignored in the neuroscience space in favor of targets specific to protein misfolding, RNA toxicity, or chemical messaging in the brain. As such, a kinase-targeting drug has not yet been approved to treat a neurological disorder. More than 15 human protein kinases have been described as phosphorylating tau, contributing to its stabilization and aggregation through phosphorylation, while others, including PDK1, PERK and GCN2, regulate amyloid precursor protein processing and protein synthesis.^*24*, *25*^ LRRK2 and PINK1 are two kinases that are mutated in PD and have garnered attention as putative PD drug targets.^*26*^ Recent disclosures of TBK1 and NEK1 as genetically-associated ALS kinases highlight the important role of kinases in ALS etiology.^*27*, *28*^ Furthermore, rapamycin, a kinase-targeting drug that enhances autophagy, was advanced to phase II for treatment of ALS.^*29*^

Driven by increased knowledge surrounding the brain penetration of kinase inhibitors, attempts have been made to repurpose FDA-approved kinase inhibitors for neurodegenerative diseases, but this strategy has not been successful.^*30*^ The majority of CNS-penetrant kinase inhibitors were optimized for systemic exposure rather than for CNS applications.^*31*^ Examples of well-studied kinases being pursued through drug repurposing are ABL1 for AD and PD (Nilotinib, Saracatinib, K0706, and Bosutinib), GSK3B for AD and PD (Tideglusib and Neu-120), MAPK14 for AD (Neflamapimod), KIT for AD and ALS (Masitinib), MLK1–3 for PD (CEP-1347), and ROCK2 for ALS (Fasudil).^*31*–*33*^ These drugs may be acting through their primary target or an off-target kinase. Saracatinib, for example, has been pursued in the case of AD due to its off-target FYN inhibition rather than ABL1 inhibition.^*34*^ Only two drugs, Masitinib for AD and ALS, and CEP-1347 for PD, made it as far as Phase III, while the rest stalled or are still being investigated (https://clinicaltrials.gov).

Many kinase-targeting small molecule drugs approved for oncology are known to potently inhibit several kinase targets and exhibit poor kinome-wide selectivity. High-quality chemical tools (potent, cell-active, narrow spectrum of kinase activity) targeting human kinases are not easy to design or identify. Given the high homology among binding sites, human kinase inhibitors require years of expert medicinal chemistry to optimize in terms of potency and selectivity. Further complications arise from the abundance of poorly characterized inhibitors in the literature and commercially available compounds that falsely advertise their quality. High-quality chemical probes offer dynamic, reversible and tunable perturbations of biomolecular functions or interactions and, importantly, serve as potential leads for drug development. While a high-quality chemical probe will require additional optimization to be used as an effective CNS-targeting agent, these small molecules represent impactful tools that can be used to validate novel targets and disease models as well as to characterize disease-propagating pathways for the first time.

## 4. A SUBSET OF UNDERSTUDIED KINASES LINKED TO NEURODEGENERATION

The majority of kinase-based research has historically focused on a subset of the human kinome, resulting in ^~^80% of kinases being overlooked.^*35*^ We refer to this portion of the kinome as understudied. While there are many ways to characterize the understudied nature of a kinase, we rely on the number of publications as one key measure to define this subset. For the purposes of this review, an understudied kinase has <66 PubMed citations on the human form of the kinase as of December 2019. This publication count was generated using the PubMed Gene database and searching for papers that are associated only with the human form of each kinase. The IDG program also uses publication count as one method to define their dark kinase list. PubMed citation count allowed us to characterize a set of 245 kinases (nearly half of the kinome) as understudied, 137 of which are also dark IDG kinases (Figure 1). We will continue to note whether a kinase is on the IDG list throughout the review.

**Figure 1.**
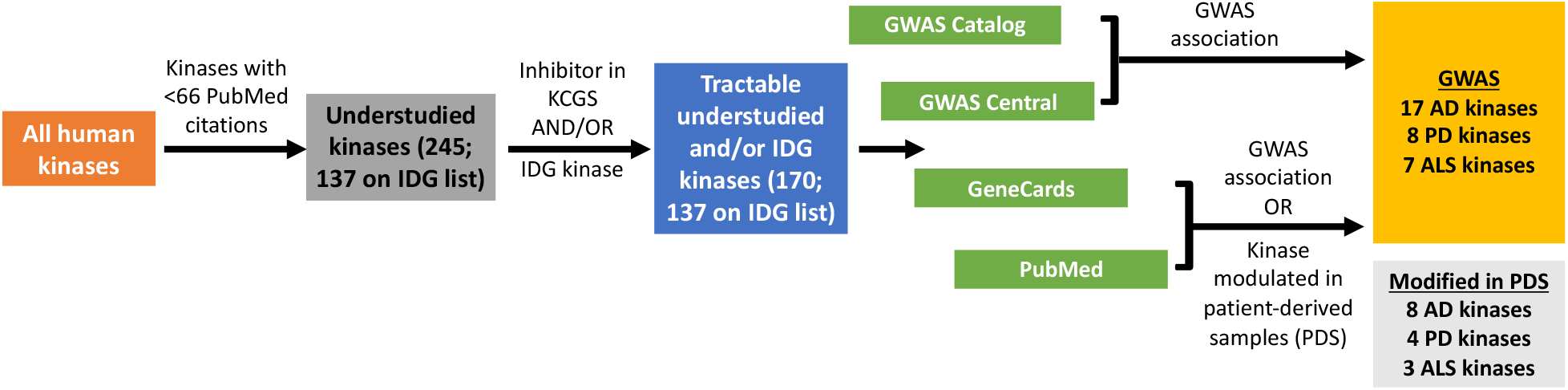
Workflow 1: Analysis of understudied kinases implicated in neurodegeneration.

This list of 245 kinases was then reduced by considering whether or not a kinase is currently tractable for drug discovery (known to bind a small molecule and potential for selectivity). Next in our analysis we checked to see if a narrow spectrum inhibitor was identified in the assembly of the kinase chemogenomic set (KCGS). KCGS was curated after a thorough examination of the data in the literature as well as for the inhibitor sets at SGC-UNC.^*36*^ If an inhibitor was not identified via this exercise then we assume that currently only promiscuous inhibitors, if any, bind that particular kinase. Of note, an inhibitor that could not be shared without restrictions was also excluded from KCGS. A final consideration we applied for inclusion of kinases in this analysis is whether the kinases are screenable. To be considered screenable for our purposes, a commercial assay must be available to profile compounds for binding and/or inhibition. As an exception to this rule, all IDG kinases were kept on the list for further consideration. This step removed 75 kinases from consideration, leaving 170 understudied kinases and preserving the 137 IDG kinases that were in the original 245 on the understudied list (Figure 1).

The understudied kinome list was further refined through text mining to determine whether each was genetically implicated in AD, PD and/or ALS. Data derived from genome wide association study (GWAS) was prioritized, followed by any data supporting mutation of a kinase and/or change in its expression in patient-derived samples (PDS). We are taking advantage of recent clinical development advances and rapid growth in GWAS datasets, partially due to advances in genomics and genetics. A recent study highlighted that drug targets with genetic support are twice as likely to be approved.^*37*^ Two databases were searched to identify relevant GWAS: https://www.ebi.ac.uk/gwas/ and https://www.gwascentral.org/. Next, the GeneCards human gene database was searched for all proteins tied to AD, PD, or ALS (https://www.genecards.org/). Any additional GWAS link that had not surfaced through searching the other two databases was noted and all publications associated with each kinase were reviewed, looking for GWAS first and patient-derived data next. Finally, a PubMed search was executed to identify any previously unidentified ties between each respective kinase and AD, PD, or ALS. GWAS were not extensively reviewed to determine the statistical significance of the implicated kinase, as an analysis of this type is beyond the scope of this review. The results of this analysis are captured in Figure 1, and in total 41 unique understudied kinases were linked to AD, PD and/or ALS through this target triage methodology. Of note, 31 of these 41 unique kinases are on the IDG list. Figure 1 captures the endpoint of *Workflow 1:* a list of kinases implicated in disease based only on text mining. Additional workflows later in the review expand on and/or refine this output.

The consequences observed in patients due to changes in expression, increased copy number or epigenetic modification, overactivation or a single nucleotide polymorphism (SNP) have been collected and published. The specific disease-propagating clinical phenotypes are summarized in Tables 1–3 along with the literature reference from which it has been abstracted. While the clinical outcome that results from modulation of each kinase is captured, the specific mechanism(s) and pathway(s) by which it drives disease are often unknown and demand additional characterization if we hope to intervene therapeutically.

**Table 1.** AD-linked kinase variants and disease implications.

| Kinase | Variant | Consequence | PubMed ID | IDG kinase |
| --- | --- | --- | --- | --- |
| ALPK2 | SNP | Increased susceptibility | 30617256 | Y |
| ALPK3 | SNP | Unknown | 22005931 | Y |
| BCKDK | SNP | Increased susceptibility | 29777097 | Y |
| CAMK1D | SNP | Increased susceptibility of LOAD | 25649863 | Y |
| CDK17 | Overexpression | Promoted AD neurodegeneration | 26885753 | Y |
| CDK18 | Overexpression | Promoted AD neurodegeneration | 26885753 | Y |
| EPHA6 | Copy number gain | Increased susceptibility of EO-FAD | 23752245 | N |
| INSRR | SNP | Increased susceptibility of LOAD | 30386171 | N |
| LMTK2 | Reduced expression | Reduced axonal transport | 31068217 | Y |
| LRRK1 | SNP | Reduced abstract reasoning | 17903297 | Y |
| MAPKAPK5 | Overexpression | Faster cognitive decline | 26080319 | N |
| MARK1 | Overexpression | Tau phosphorylation | 11089574 | Y |
| MARK4 | SNP | Increased susceptibility | 30617256; 30979435 | Y |
| MAST4 | SNP | Reduced hippocampal volume | 28098162 | Y |
| MINK1 | SNP | Increased susceptibility | 30413934 | N |
| NEK10 | SNP | Increased susceptibility of LOAD | 27770636 | Y |
| NUAK1 | SNP | Increased cortical Aβ levels | 23419831 | N |
| PIM3 | Overexpression | Increased susceptibility | 30349106 | N |
| SIK1 | SNP | Increased susceptibility of FAD | 26830138 | N |
| STK32B | SNP | Faster cognitive decline | 23535033; 30636644 | Y |
| STK32C | Increased methylation | Accelerated Braak stage progression | 30045751 | Y |
| TAOK1 | Overexpression | Tau phosphorylation | 29730992; 23585562 | Y |
| TAOK2 | Overexpression | Tau phosphorylation | 29730992; 23585562 | Y |
| TNK1 | SNP | Increased susceptibility | 28631188; 18830724 | N |
| TTBK1 | SNP | Increased susceptibility of LOAD | 21219968 | Y |

LOAD: late-onset AD; EO-FAD: early-onset familial AD; FAD: familial AD

**Table 2.**
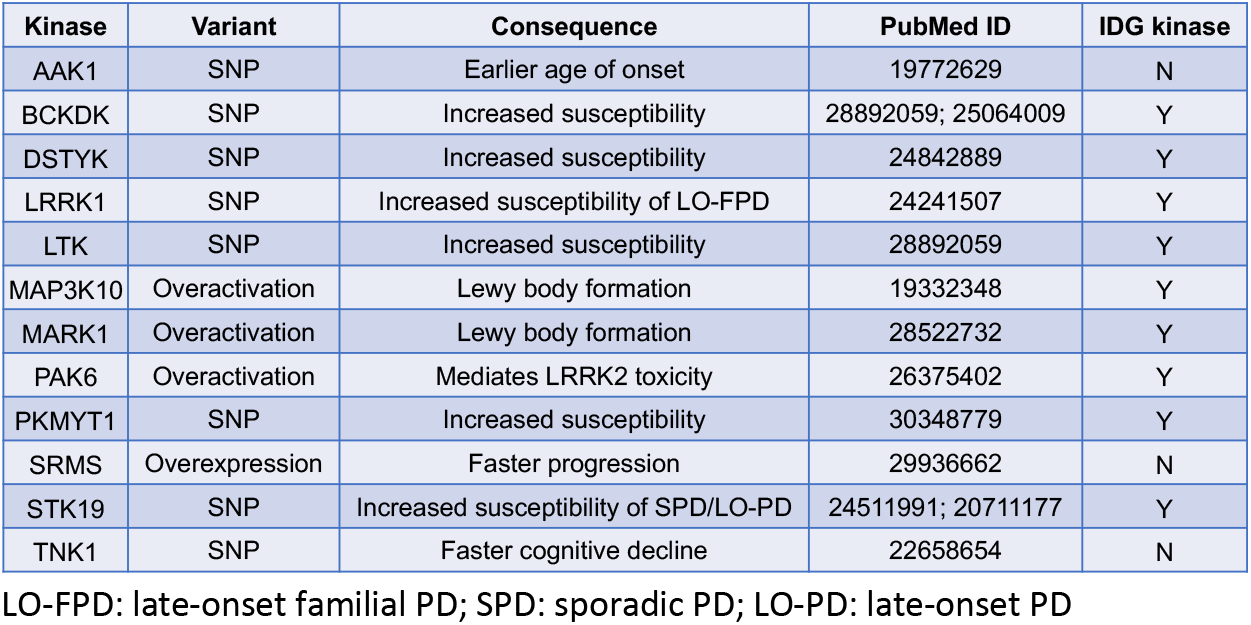
PD-linked kinase variants and disease implications.

**Table 3.** ALS-linked kinase variants and disease implications.

| Kinase | Variant | Consequence | PubMed ID | IDG kinase |
| --- | --- | --- | --- | --- |
| AAK1 | Decreased expression | Disregulated vesicle recycling | 25514244 | N |
| CAMK1G | SNP | Increased susceptibility | 23624525 | Y |
| CSNK1G3 | SNP | Increased survival | 19451621 | Y |
| DYRK2 | SNP | C9orf72 mutation interaction | 22959728 | Y |
| NEK1 | SNP | Increased susceptibility | 25700176 | Y |
| PXK | SNP | Longer telomeres; longer survival | 30931641 | Y |
| SCYL3 | SNP | Increased survival | 19451621 | Y |
| STK36 | SNP | Increased association with ALS | 22959728 | Y |
| TTBK1 | Overexpression | TDP-43 phosphorylation | 25473830 | Y |
| TTBK2 | Overexpression | TDP-43 phosphorylation | 25473830 | Y |

## 5. KINASE EXPRESSION ANALYSES

An interesting cross-examination is to compare publication count and expression of both understudied as well as well-studied kinases in a disease-relevant cell type. While expression is not the only determinant as to whether a kinase is significant in a disease, the complete absence of said kinase in a specific cell population diminishes its probable impact in driving disease pathology. The subset of understudied kinases defined for each respective disease as well as 5 well-studied kinases (Table 4) was plotted versus publication count (blue bars) and RNA expression (orange lines) for AD, PD, and ALS (Figures 3–5). Kinases are oriented along the x-axis in order of increasing number of publications. As described previously, the publication count as of December 2019 was tabulated via searching for papers associated with the human form of each kinase using the PubMed Gene database.

**Table 4.**
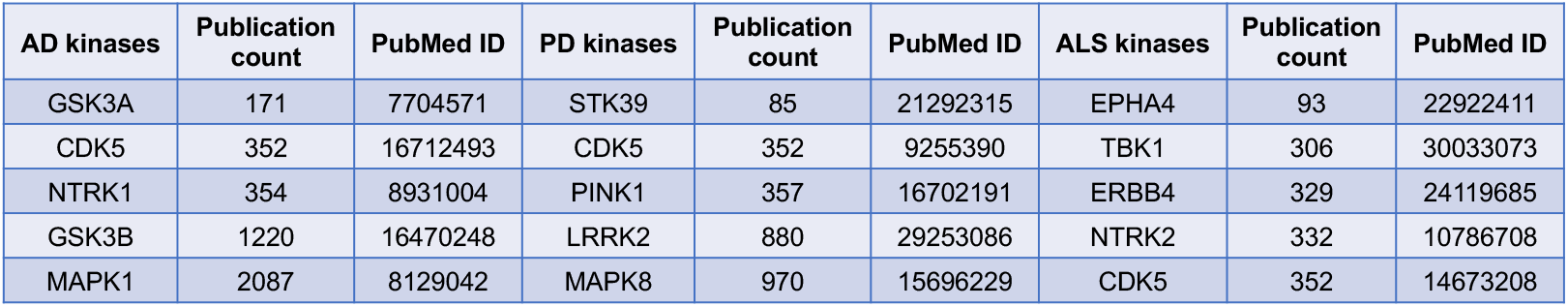
Examples of well-studied kinases in AD, PD, and ALS.

Using the GEO DataSets repository (https://www.ncbi.nlm.nih.gov/gds/), we analyzed the specific brain regions and cell types that are impacted in chronic neurodegenerative diseases for their RNA expression of kinases identified via *Workflow 1* (Figure 1). *Workflow 2* (Figure 2) outlines the method by which we curated the kinase list generated from *Workflow 1* via analyzing expression of these disease-associated kinases in disease-relevant cells. For AD, we opted to examine and summarize kinase expression in microglia derived from human induced pluripotent stem cells (iPSCs) donated by healthy volunteers and published as a bulk RNA sequencing dataset (GSE133432, Figure 3).^*38*^ For PD, we analyzed RNA sequencing data collected from human midbrain DA neurons from the ventral tegmental area and SN pars compacta of human post-mortem brains (GSE76514, Figure 4).^*39*^ For ALS, human single MNs from healthy volunteers were sequenced and RNA data plotted (GSE121069, Figure 5).^*40*^

**Figure 2.**
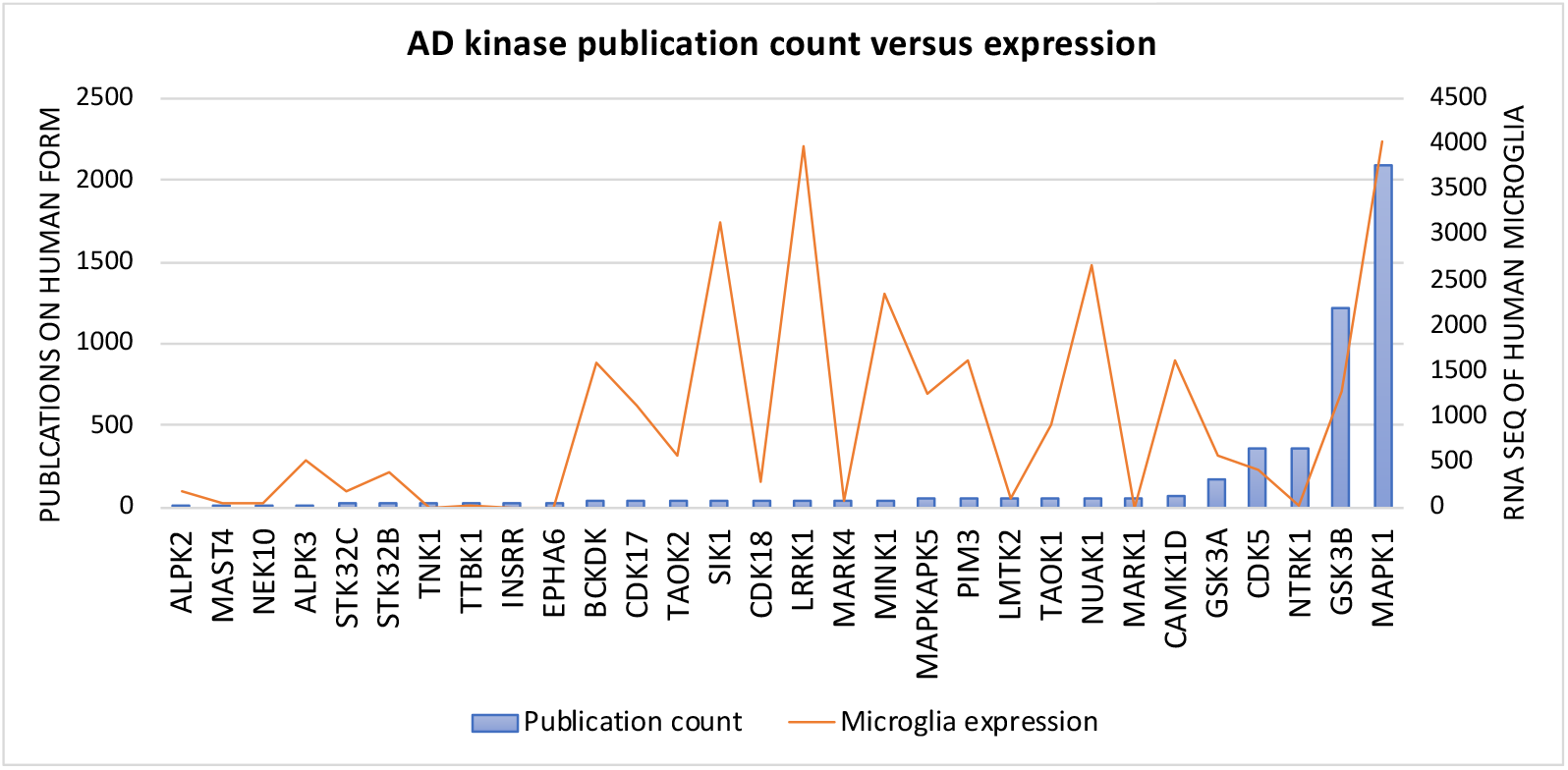
Publication count versus expression data for kinases implicated in AD.

**Figure 3.**
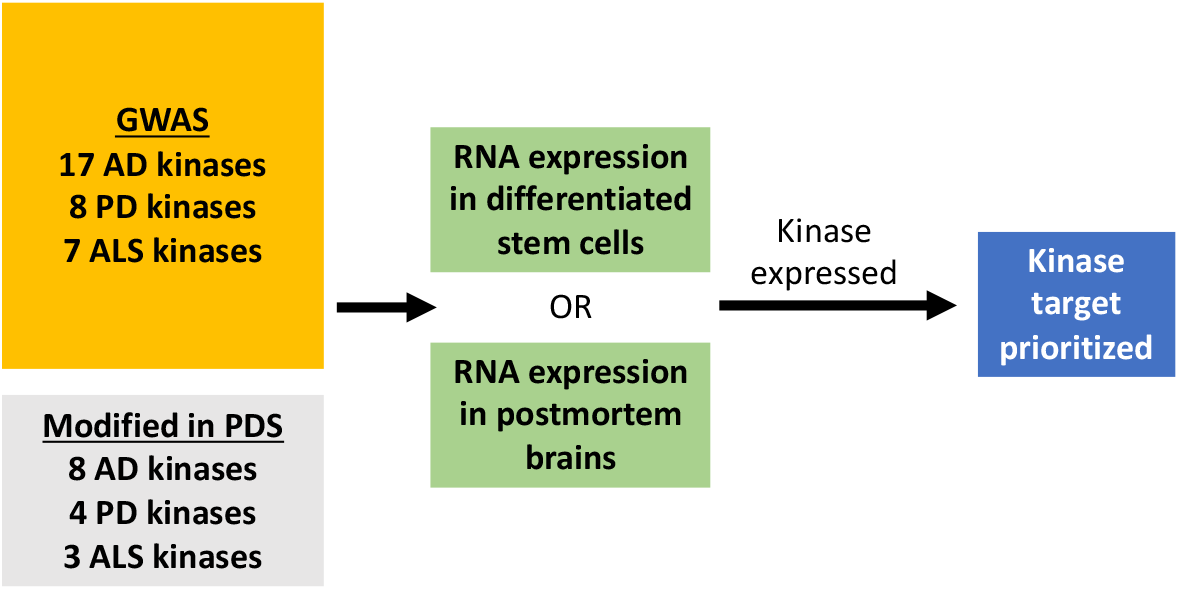
Workflow 2: Curation of kinases via expression analyses.

**Figure 4.**
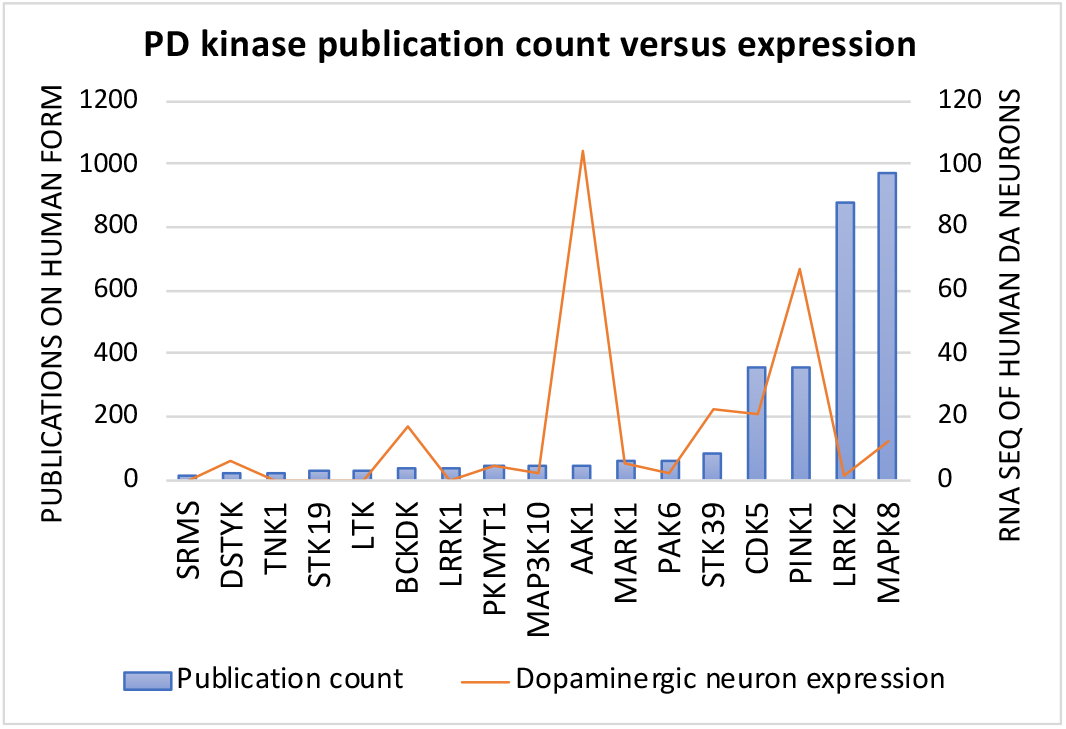
Publication count versus expression data for kinases implicated in PD.

**Figure 5.**
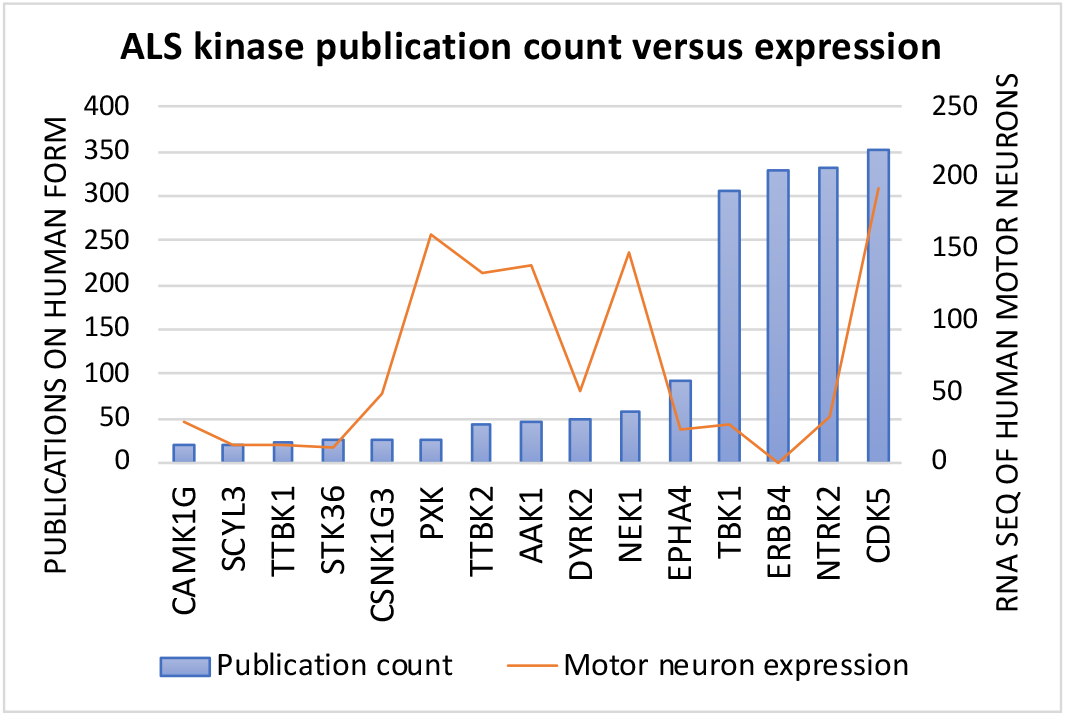
Publication count versus expression data for kinases implicated in ALS.

Included in our study are some of the more studied kinases implicated in these same diseases. These kinases were identified using the GeneCards database. All proteins associated with a disease are given a significance score by the GeneCards website that roughly corresponds with how relevant and validated they are with respect to that specific disease. We selected the top five kinases on the list corresponding to each disease ranked with the highest significance scores (Table 4). The publication count as of December 2019 obtained through tabulating papers that reference the human kinase using the PubMed Gene database as well as the relevant representative publication linking the kinase to disease are also provided in Table 4. None of these well-studied kinases are on the IDG list.

Interestingly, examining the relationship between publication count (blue bars) and RNA expression (orange lines) as shown in Figures 3–5 clearly demonstrates that those kinases which have been focused on by the neurodegeneration field to date are not necessarily the most relevant from an expression standpoint. *Workflow 2* (Figure 2) provides a method by which kinases implicated through mining patient genetic data can be refined prior to the experimental phase of neuroscience-focused kinase drug discovery.

## 6. NEW TARGETS AND PRE-CLINICAL CANDIDATES FOR AD, PD, AND ALS

This section focuses on kinases that are understudied, implicated in at least one chronic neurodegenerative disease, expressed in relevant cell type(s), and for which a narrow spectrum inhibitor has been identified. While patient-derived data has implicated these kinases in one or more neurodegenerative disorder, often more characterization has to be done to determine whether inhibition will elicit a positive outcome for patients. The lack of research on these kinases has also left many of them with few good chemical tools to study their biology, making KCGS an invaluable resource to learn more about their roles in neurodegeneration.

We have selected a subset of these kinases to highlight. *Workflow 3* (Figure 6) outlines how we identify kinase inhibitors and the next steps once these inhibitors are selected. There are public databases and publications that harbor kinase screening data from which we can identify potential inhibitors and start to assess their selectivity/potency. If a kinase has not been screened previously, development of an assay that enables identification of inhibitors would be the first step in *Workflow 3*. The best available inhibitor(s) in terms of potency and selectivity across the kinome can then be advanced into a disease-relevant assay to begin to determine if (1) the target is valid and (2) the compound can be considered a lead in the drug discovery pipeline. Selected output from *Workflow 3* is captured in Table 5. Compounds include those from KCGS, the best available inhibitors that have been published but are not included in KCGS, and IDG DK (dark kinase) tools. As part of the IDG program, we confirm cellular penetration and potent (<1 μM) target engagement of the kinase target before naming a compound a DK tool. The understudied nature of these kinases has resulted in only a few available chemical tools to further characterize their role in neurodegeneration. While these small molecules are not optimized for CNS applications, they represent a starting point for interrogating biological pathways that are aberrant in AD, PD, and/or ALS. Structures corresponding to compounds listed in Table 5 are shown in Figure 7 with their hinge-binding moieties oriented down in all cases. In accordance with our aspirational probe criteria,^*41*^ kinome-wide selectivity needs to be improved and activity in cells needs to be confirmed before any one of the molecules in Table 5 can be named a chemical probe for the target of interest. Important future work for the field is implementation of iterative medicinal chemistry optimization efforts to convert these narrow spectrum starting points into high-quality chemical probes. With high-quality probes available, the community will be able to interrogate the impact of inhibition in brain disease signaling. When moving beyond cell-based assays into animal models and eventually humans, blood–brain barrier penetrance will have to be assessed and improved via medicinal chemistry.

**Figure 6.**
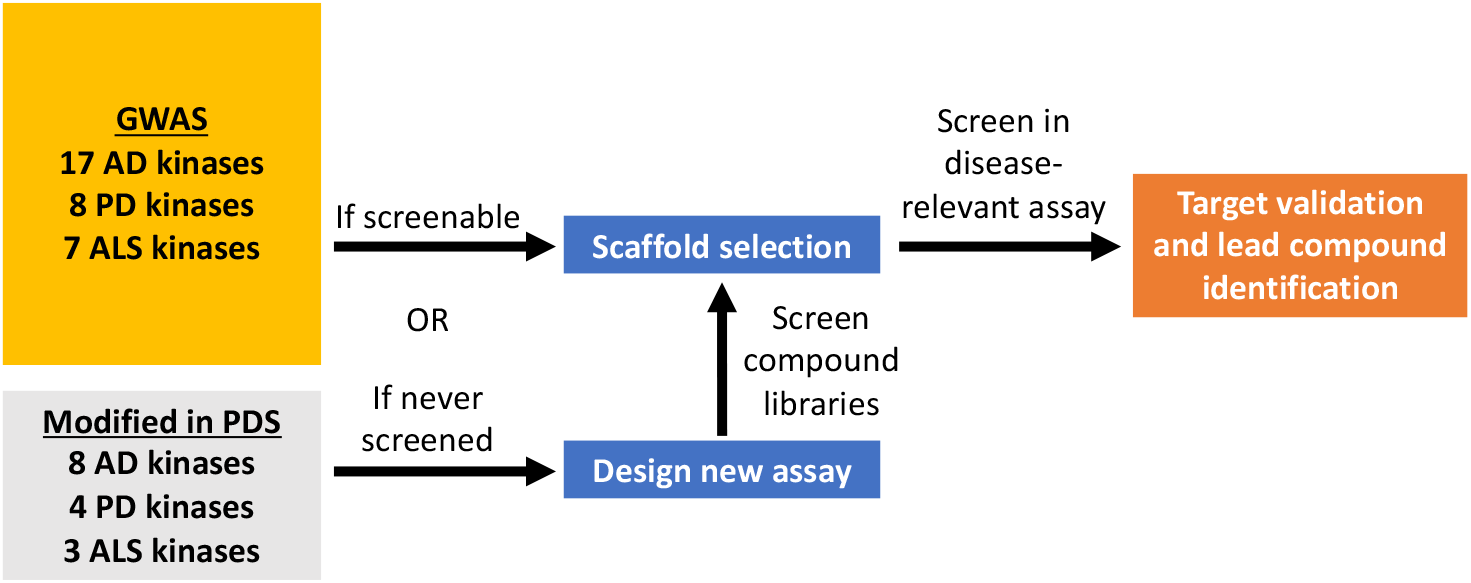
Workflow 3: Steps to validate kinase target and identify a small molecule to advance.

**Figure 7.**
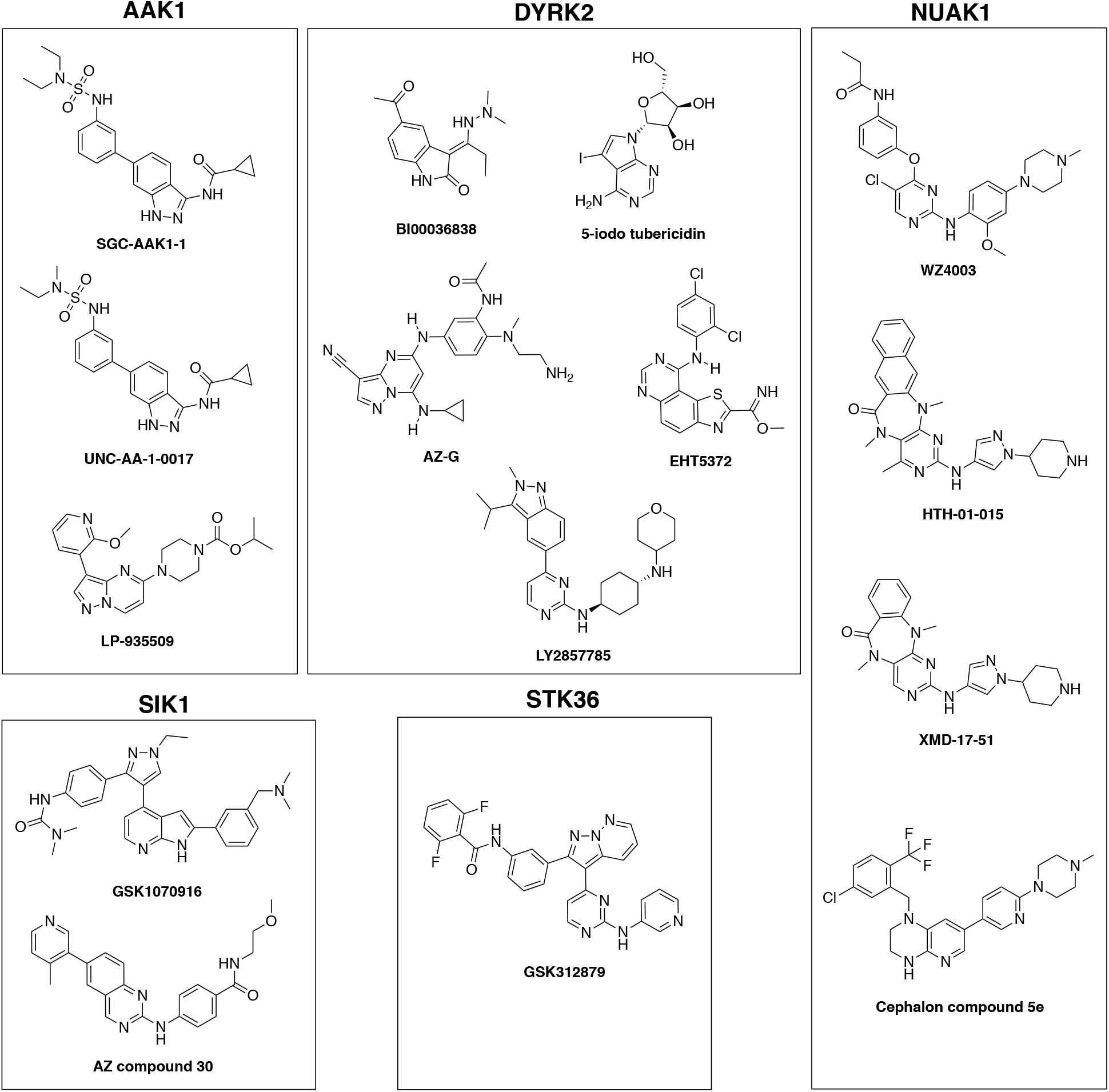
Structures corresponding to narrow spectrum inhibitors of kinases listed in Table 5.

**Table 5.** Molecules that inhibit understudied kinases with AD, PD, and/or ALS implications.

| Kinase | AD, PD, ALS link | KCGS coverage | Literature compounds | DK tools |
| --- | --- | --- | --- | --- |
| AAK1 | GWAS PD and ALS PDS | SGC-AAK1-1<sup>42, 43</sup>, UNC-AA-1-0017<sup>43</sup> | LP935509<sup>44</sup> | Not IDG |
| STK36 | GWAS ALS | GSK312879<sup>45</sup> | - | - |
| SIK1 | GWAS AD | GSK1070916<sup>46</sup> | AZ compound 30<sup>47</sup> | Not IDG |
| DYRK2 | GWAS ALS | BI00036838<sup>48</sup> | EHT5372,<sup>49, 50</sup> LY2857785,<sup>51</sup> 5-iodo tubericidin<sup>52</sup> | AZ-G<sup>53</sup> |
| NUAK1 | GWAS AD | HTH-01-015, WZ4003, XMD-17-51<sup>54</sup> | Cephalon compound 5e<sup>55</sup> | Not IDG |

AAK1 is inhibited by the 3-acylaminoindazole scaffold in KCGS. SGC-AAK-1, in fact, was recently identified as a selective chemical probe for studying AAK1 biology and released alongside a structurally related negative control compound that does not inhibit AAK1.^*42*, *43*^ AAK1 expression is increased in PD patient brains and was correlated with age of onset in a GWAS (Table 2).^*56*, *57*^ The influence of AAK1 was suggested to rely on its essential role in endocytosis and lysosomal sorting, which, when aberrant, can contribute to the pathology of PD.^*56*^ AAK1 was shown to impact alpha-synuclein aggregation and resultant neurodegeneration in *C. elegans*, and its inhibition has been proposed to be therapeutically advantageous in the case of PD.^*58*, *59*^ In contrast, AAK1 protein levels were found to be decreased in ALS patients (Table 3). Furthermore, AAK1 was found to interact with mutant but not wild-type SOD1 in a yeast two-hybrid system and misdirected into aggregates containing mutant SOD1 and neurofilament proteins in rodent models of ALS. Dysfunction of AAK1 drives ALS pathology since its role is essential in the endosomal and synaptic vesicle recycling pathway.^*60*^ Further characterization of AAK1 with respect to ALS will determine whether inhibition is favorable in interrupting disease progression.

STK36 is an IDG kinase inhibited by a single compound in KCGS that was donated by GlaxoSmithKline. STK36 regulates the activity of GLI zinc-finger transcription factors, which impacts Hedgehog signaling, cell proliferation, and cell-fate specification.^*61*^ STK36 was a notable locus that showed increased evidence of association with ALS across 13 patient cohorts (Table 3).^*62*^ Given its role in regulating transcription, STK36 variants are thought to contribute to abnormal RNA transcription and processing in ALS.^*63*^ The impact of STK36 inhibition in the case of ALS can be interrogated further using the small molecule that we have described.

SIK1 is an understudied kinase inhibited by a single molecule in KCGS. A number of studies in the last decade have implicated SIK1 in brain pathology.^*64*^ A GWAS examining genes that contribute to AD age of onset identified SIK1 as a gene in close proximity to a SNP (Table 1).^*65*^ SIK1 did not achieve genomewide significance in this study. Importantly, significantly increased expression of SIK1 was observed in neutrophils taken from AD patients demonstrating dementia.^*66*^ It has been proposed that inhibition of SIK1 in pre-clinical mouse models of AD results in reduced neuroinflammation.^*67*^ This finding is in alignment with the determination that SIK1 is crucial to regulating microglial apoptosis, especially given that microglial activation leads to neuroinflammation and is a hallmark of many neurodegenerative processes.^*64*^ Since neuroinflammation exacerbates AD, possibly leading to nerve cell death and synaptic dysfunction, inhibition of SIK1 would offer an attractive method to reduce this pathology-driving process.

DYRK2 is an IDG kinase inhibited by a single molecule in KCGS. We prepared a small library of pyrazolopyrimidines based upon a literature compound that was reported to have DYRK2 activity.^*53*^ We profiled this library using the cell-based DYRK2 nanoBRET assay^*68*^ and we were able to identify AZ-G as meeting the criteria to be named a DK tool for DYRK2 (IC_50_ = 160 nM). Details about the DYRK2 nanoBRET assay and DK tool can be found at https://darkkinome.org/kinase/DYRK2. DYRKs are a family of conserved kinases that play key roles in the regulation of cell differentiation, proliferation, and survival.^*69*^ DYRK2 has been associated with ALS via GWAS. A genomic region that correlates with DYRK2 was identified as impacting the age of onset of ALS (Table 3).^*62*^ Additional studies are required to determine whether pharmacological inhibition of DYRK2 is advantageous in the case of ALS. DYRK2 has also been shown to phosphorylate tau i*n vitro* on a residue that is hyperphosphorylated in filamentous tau from AD brains.^*70*^ This suggests that inhibition of DYRK2 may be therapeutically beneficial for AD.

NUAK1 is inhibited by three KCGS molecules. This kinase is highly expressed early in the developing brain.^*71*^ NUAK1 was reported as a biomarker for AD in a GWAS (Table 1). Through assessment of Aβ levels *in vivo* via PET imaging, NUAK1 was associated with quantitative global Aβ loading.^*72*^ Furthermore, NUAK1 was found in parallel cell-based and Drosophila genetic screens to regulate tau levels. Inhibition of NUAK1 in fruit flies suppressed neurodegeneration in tau-expressing Drosophila and NUAK1 haploinsufficiency rescued the phenotypes of a tauopathy mouse model.^*73*^ These cell and animal models point to NUAK1 inhibition as a potentially beneficial strategy in treating AD.

## 7. PATH FORWARD

We have identified kinases that are implicated in AD, PD, and/or ALS due to mutations, differential epigenetic processing, enhanced activation, or altered expression in patients. The clinical outcome of these changes, such as earlier onset of disease or prolonged life span, has been established in most cases (Tables 1–3). Relevant data that is missing is how modulating the kinase impacts disease-propagating pathways. Studying neurodegenerative disorders has proved to be challenging in the past due to the inaccessibility of human cells from individuals affected by the disease, with post-mortem tissue only providing a snapshot of the final stage of the disease. While immortalized cells lines and animal models have greatly increased our understanding of key biochemical pathways implicated in the pathogenesis of the disease, they fail to truly model the disease observed in humans as there are considerable differences between cell lines and between species in terms of protein expression levels, signaling pathways, and cellular processes.^*74*–*76*^ This gap in translation may be one of many reasons why hundreds of clinical trials for neurodegenerative treatments have failed in the past couple of decades.^*18*, *22*, *77*^ iPSCs offer the capability to generate specific patient-derived neural subtypes, with CRISPR-Cas9 technology allowing for the genetic modification of these cells.^*78*^ Since neurodegenerative diseases are highly complex, with many different cell types and regions involved, 3D cultures are a step forward in attempting to reproduce the multidimensional environment of the brain and hold an immense potential for modeling neurodegenerative diseases.^*74*^ Figure 8 highlights the entire process from patient to drug, highlighting the enabling technologies of iPSCs combined with CRISPR-Cas9 to both model disease and discover/test new therapeutics.

**Figure 8.**
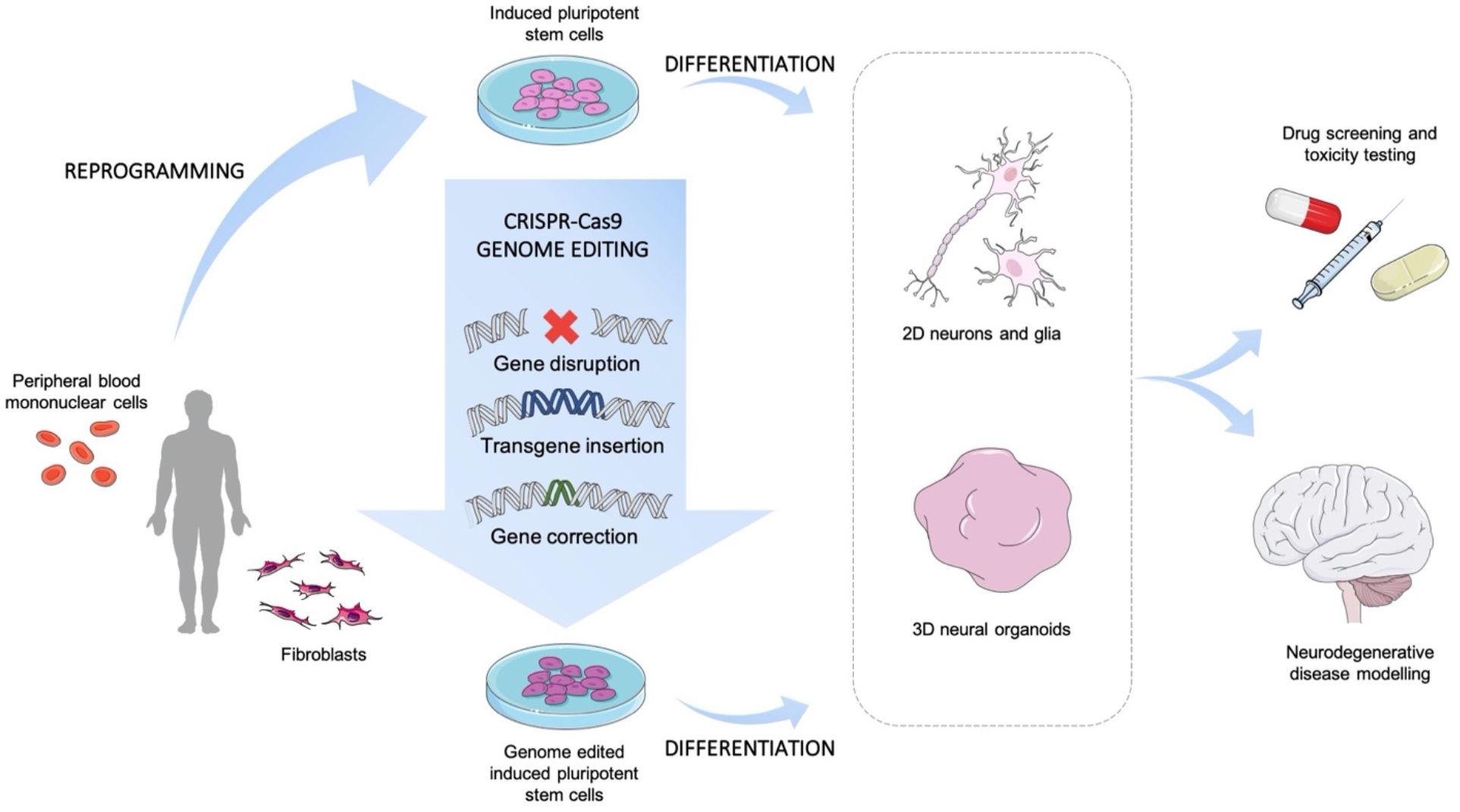
Human iPSCs as tools for drug screening and neurodegenerative disease modeling.

This system can be used to determine the results of kinase knockout or acute inhibition and bridge the gap between a novel kinase target and observed patient phenotype. The phenotype that results from acutely inhibiting one of these kinases may differ from what is observed due to chronic genetic knockout. Recent studies have revealed that genetic compensation in response to gene knockout can explain the different results obtained by acute and chronic gene inhibition.^*79*, *80*^ Importantly, studies of this type will help determine whether inhibition is a therapeutically beneficial strategy for each kinase tested.

An aspirational goal is to develop small molecules for all kinases that have interesting biology related to AD, PD, and/or ALS but for which good chemical starting points do not currently exist or are in very early stages of development. Here we share some examples of kinases in this category that deserve medicinal chemistry lead discovery effort. NEK1 is an IDG kinase in need of a good chemical tool to enable its study.^*81*^ NEK1 variants confer susceptibility to ALS. Significant association was identified between loss-of-function NEK1 variants and familial ALS risk (Table 3).^*28*^ The fact that loss-of-function is causative makes NEK1 a special case, where restoration rather than inhibition of its cellular role is required. TTBK1 and TTBK2 are two CK1 family kinases that are both IDG targets without good inhibitors described. SNPs in the TTBK1 gene have been associated with late-onset AD. Specific polymorphisms were found to decrease the risk of late-onset AD in multiple patient populations (Table 1).^*82*^ Abundant expression of TTBK1 in the cortical and hippocampal neurons of AD brains, coupled with its confirmed role of phosphorylating tau and leading to pre-tangles, suggests that TTBK1 inhibitors would be therapeutically beneficial to AD patients.^*83*^ Furthermore, both TTBK1 and TTBK2 co-localize with TDP-43 inclusions in the ALS spinal cord and, thus, could represent attractive targets for therapeutic intervention for ALS (Table 3).^*84*^

TAOK1 and TAOK2 are understudied IDG kinases. Good molecules that inhibit the TAOKs are needed. TAOK1 and TAOK2 are phosphorylated and active in AD brain sections and they co-localize with both pre-tangle and tangle structures. TAOK inhibition was found to reduce tau phosphorylation in vitro and in cell models.^*85*^ Finally, STK19 is an IDG kinase that has been genetically implicated in PD via two separate GWAS but suffers from a lack of chemical inhibitors. The gene coding for STK19 was flagged as a highly significant locus with a strong association in sporadic PD (Table 2).^*86*^ An siRNA library screen highlighted that STK19 regulates levels of phosphorylated alpha-synuclein, implicating a role in the development of Lewy bodies and the pathology of PD.^*87*^ Further supporting its function in the brain, STK19 was also detected as a susceptibility gene for schizophrenia via GWAS.^*88*^ Dedicated medicinal chemistry efforts are required to develop good chemical tools to study these kinases.

As part of our efforts to expand KCGS and increase its kinome-wide coverage in subsequent releases, we will develop narrow spectrum compounds that inhibit some of the neurodegeneration-implicated kinases discussed in this review that currently lack chemical coverage. We will also target several of these kinases through the IDG program, delivering cell-active DK tools to facilitate their biological characterization. Kinase inhibitors have proven useful for elucidation of the roles of particular kinases in signaling pathways and in the role these pathways play in disease. Importantly, this knowledge has led to more than 50 FDA-approved medicines that help patients. In spite of this success, it is surprising that much of the kinome remains unexplored. The lack of good chemical tools limits our ability to realize the value of understudied kinases. This is especially true for the neural kinome. We have highlighted the potential of targeting kinases expressed in the brain and implicated in neurodegeneration for the treatment of AD, PD, and ALS, and provided workflow that could offer insight into their biology. A concerted community-wide effort is required to illuminate the impact of these understudied neural kinases on disease-propagating pathways in the brain. Illumination will most effectively be realized if research is done collaboratively and in the open, sharing our findings in real time with the research community at large.

## AUTHOR INFORMATION

### Author Contributions

A.D.A. and D.H.D. conceived the idea. A.D.A conducted literature review and drafted the manuscript. D.H.D., C.W., and A.D.A. identified promising kinase inhibitor scaffolds. A.I.K. and T.D. contributed to writing the manuscript. All authors contributed to critical editing and preparation of the final draft. All authors read and approved the final manuscript.

### Funding

The Structural Genomics Consortium is a registered charity (number 1097737) that receives funds from AbbVie, Bayer Pharma AG, Boehringer Ingelheim, Canada Foundation for Innovation, Eshelman Institute for Innovation, Genome Canada, Genentech, Innovative Medicines Initiative (EU/EFPIA) [ULTRA-DD grant no. 115766], Janssen, Merck KGaA Darmstadt Germany, MSD, Novartis Pharma AG, Ontario Ministry of Economic Development and Innovation, Pfizer, São Paulo Research Foundation-FAPESP [2013/50724-5, 2014/5087-0 and 2016/17469-0], Takeda, and Wellcome [106169/ZZ14/Z]. A.D.A., D.H.D. and C.W. acknowledge U24DK116204 for support. A.D.A. and C.W. acknowledge ALSA Investigator Initiated Starter Grant wa1127 and U54AG065187 for support. T.M.D acknowledges the Ghislaine and Sebastian Van Berkom foundation and McGill Healthy Brains for Healthy Lives for funding support.

### Notes

The authors declare no competing financial interest.

## ABBREVIATIONS

CNS: central nervous system
KCGS: kinase chemogenomic set
FDA: Food and Drug Administration
AD: Alzheimer’s disease
SGC: Structural Genomics Consortium
PD: Parkinson’s disease
ALS: amyotrophic lateral sclerosis
IDG: Illuminating the Druggable Genome
Aβ: amyloid-β
NFTs: neurofibrillary tangles
DA: dopamine
SN: substantia nigra
REM: rapid eye movement
RBD: REM sleep behavior disorder
MN: motor neuron
NMDA: N-methyl-D-aspartate
L-DOPA: Levodopa
GWAS: genome wide association study
PDS: patient-derived sample
SNP: single nucleotide polymorphism
iPSCs: induced pluripotent stem cells

